# CNAHap: a germline haplotyping method using tumor allele-specific copy number alteration

**DOI:** 10.1101/2021.03.27.437314

**Authors:** Bowen Tan, Lingxi Chen, Wenlong Jia, Yanfei Wang, Hechen Li, Shuai Cheng Li

**Author notes:** These authors contributed equally to this work (Bowen Tan, Lingxi Chen).

## Abstract

Haplotype phasing is indispensable to study human genetics. The pervasiveness of large copy number variant segments in solid tumors brings possibilities to resolve long germline phasing blocks utilizing allele imbalance in tumor data. Although there exist such studies, none of them provide easy-use software based on availability and usability. Herein, we present a novel tool, CNAHap, to determine the allele-specific copy number in tumor and then phase germline variants according to the imbalanced alleles in tumor genomes. We also provide interactive web interfaces to visualize the copy number and phase landscape from CNAHap. On *in silico* datasets, CNAHap demonstrates higher allele-specific copy number calling accuracy than the benchmark tool and generates long phasing blocks. As a case study on Hepatocellular carcinoma, CNAHap successfully generates huge phase blocks with the averages of N50 and N90 as 25M and 7M, respectively, and finds the Olfactory receptor family is recurrent amplified. Our results illustrate the efficacy of CNAHap in determining tumor allele-specific copy numbers and their long germline haplotypes. CNAHap is available at https://github.com/bowentan/CNAHap and the CNAHap visualization web interfaces are hosted at bio.oviz.org.

## Introduction

The human genome consists of pairs of paternal and maternal chromosomes. The pairs of homologous chromosomes differentiate with minute genomic variations, including single-nucleotide variations (SNVs), small insertions and deletions (InDels), short tandem repeats (STRs), *etc*. (1). High-throughput sequencing protocols profile reads from a mixture of two homologous chromosomes, thereby failing to determine the chromosome origin of a sequencing read. Accordingly, for a couple of heterozygous loci whose genomic distance is farther apart than the sequencing read length and insertion size, whether the alleles are from identical chromosomes is concealed (2). Haplotype phasings reveal heterozygous SNV and InDel loci to their corresponding paternal or maternal haplotype from the sequencing observation (3). Accurate whole genome wide phasing sheds light on medical genomics (4, 5) and population genetics (6, 7).

Diverse methods exist for resolving haplotypes from wetlab methods or sequencing data. Laboratory-based phasing methods are costly or impractical due to laborious efforts (8). Current popular computational approaches for phasing haplotypes employ two strategies (9). The first one utilizes the population database to phase while demonstrates the inability of handling rare and *de novo* variants (10). The latter strategy is to assemble the haplotype from the sequencing reads. Mainstream haplotype assembly tools catalog the genetic variants of the germline haplotype by incorporate the linkage information from high-throughput sequencing of normal tissue (11–16). Nevertheness, the length of the phased block, and the number of phased SNVs/InDels rely on the read linkages.

To further extend the phased block, some studies incorporate tumor data to unveil germline haplotypes. Large somatic copy number aberration (SCNA) blocks are prevalent (almost 90%) in solid tumors (17). Equipped with tumor allele frequency, now scientists can phase over the large copy number aberration (CNA) blocks and are free from the read length and insert size of a sequencing protocol, promoting a higher phase rate than merely adopting normal data (18). HATS (19) is a population-based approach that adopts a hidden Markov model to construct germline haplotypes in copy number variation (CNV) gain regions. VAF phasing (18) forms germline haplotypes by distinguishing variant allele frequency (VAF) changes between paired tumor and normal tissues in areas of CNV gains. However, running these tools requires arduous user interventions as VAF phasing provides no open-source software and HATS necessitates a training process first.

In this work, we spotlight germline phasing with tumor CNA, and propose a novel user-friendly tool, CNAHap (https://github.com/bowentan/CNAHap, Figure 1), to phase SNVs/InDels as in normal cells by taking advantage of allele imbalance from paired tumor CNV blocks. CNAHap also calls the allele-specific copy number aberrations in tumor cells. In addition, to visualize the CNAHap output vividly, we developed three online interactive visualization applications (CNV: Circos View, CNV: Focal Cluster, and Phased: On Genes) hosted in Bio-Oviz bio.oviz.org (20) (Table 1). We validated the phasing efficacy of CNAHap in three *in silico* WGS data sets with different tumor purity rates, and CNAHap exhibits a highel allele-specific copy number calling accuracy than the benchmark tool and generates long phasing blocks. Then we conducted a case study in a Hepatocellular carcinoma (HCC) cohort. CNAHap successfully generates huge phase blocks with the average N50 and N90 as 25M and 7M, respectively, and finds the Olfactory receptor family is recurrent amplified.

**Fig. 1.**
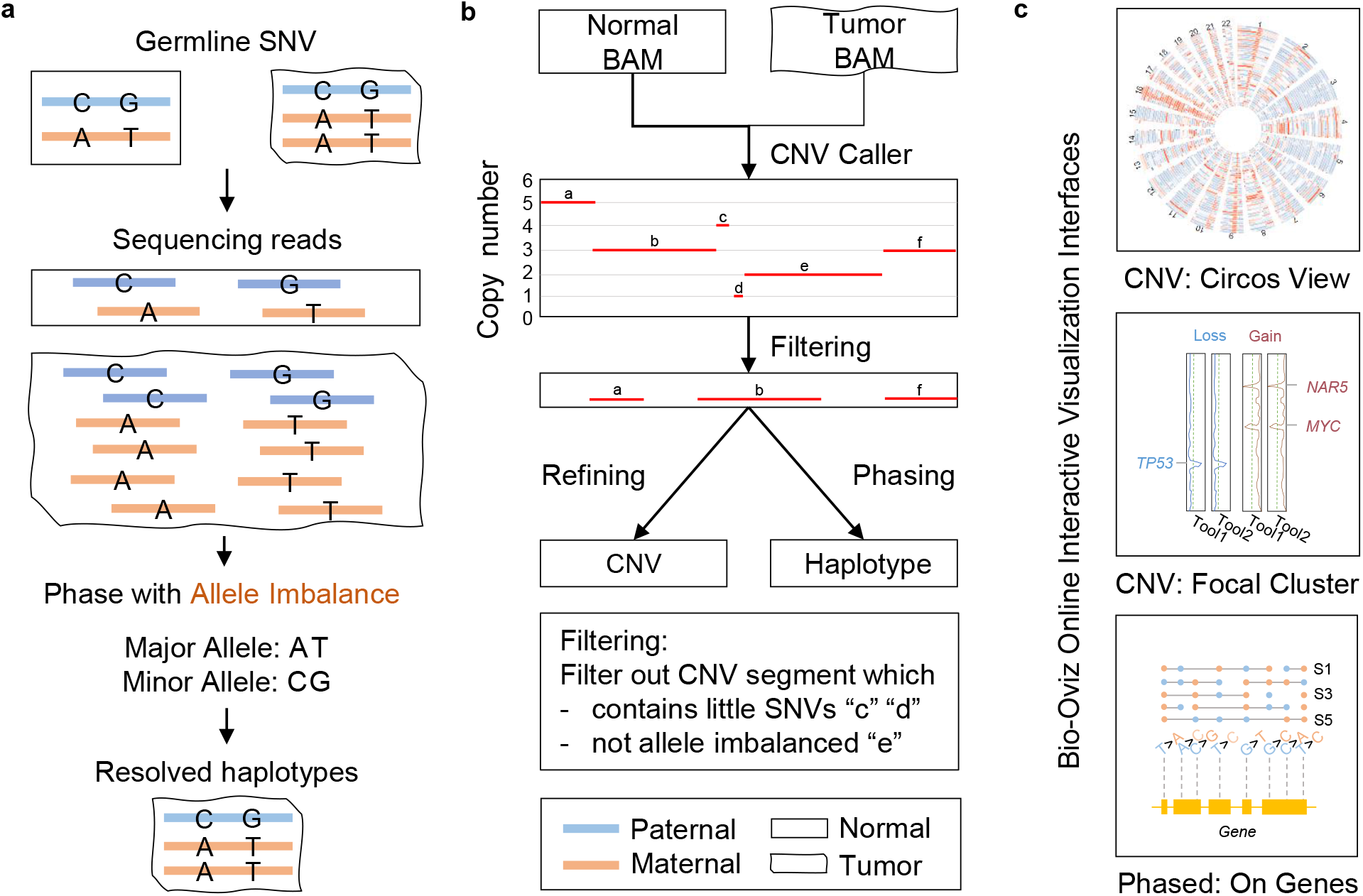
Overview of CNAHap. (a) The core principle of CNAHap to phase. (b) The workflow of CNAHap. (c) Three online interactive visualization interfaces hosted in bio.oviz.org Oviz-Bio (20).

**Table 1.** Summary of CNAHap visualization interfaces in bio.oviz.org Oviz-Bio (20).

| CNAHap visualization web interface | Key Functionalities |
| --- | --- |
| CNV: Circos View, e.g., Figure 3a<br>https://bio.oviz.org/demo-project/analyses/CNV_circos_view | Circos plot of total, major, and minor copy number<br>Circos plot of allele imbalance<br>Circos plot of phased information |
| CNV: Focal Cluster, e.g., Figure 3b<br>https://bio.oviz.org/demo-project/analyses/CNV_focal_cluster | Recurrent gains and losses<br>Gene annotation Ensembl (32)<br>Illustrates multiple tools results parallelly |
| Phased: On Genes, e.g., Figure 5, Supplementary Figure S2<br>https://bio.oviz.org/demo-project/analyses/Phased_on_genes | Phasing profile on genes, mutation detail information<br>Genes, transcripts annotation Ensembl (32)<br>Protein annotation Pfam (33) |

## Materials and methods

To estimate germline haplotypes from normal and tumor samples, CNAHap consists of two components. The first is to estimate allele-specific copy numbers, i.e., the copy numbers of the two haplotypes of given segments in tumor cells, The second is to perform the SNV phasing on segments by the fact that SNVs along the same haplotype sharing a similar copy rate. When a CNV event occurs, the alleles with larger, depths seem to be along one haplotype, and the alleles with smaller depth aligns with the other haplotype.

### Target SNV extraction

Given the sequencing data of normal and tumor cell samples from a cancer patient, CNAHap is designed to find two haplotypes of SNV loci which are supposed to be originated from the germline, hence they are contained extensively in all types of cells, such as tissue and germline cells. Therefore, a shared set of heterozygous SNVs as the target SNVs loci are extracted from normal and tumor cell samples by selecting SNVs with the same identifiers including contig names, positions, reference alleles and alternative alleles between normal and tumor cells. All subsequent analysis will be performed merely on these target SNVs.

### Allele imbalance and copy number estimation

Before estimating haplotypes, CNAHap first needs to estimate allele-specific copy numbers of the given CNV segments. If a CNV event occurs in a genomic region or a genomic segment, three possible outcomes will arise. The first is segments with imbalanced copies, because of different numbers of copies two haplotypes are duplicated. The second is balanced segments with the same number of copies. The third outcome is that one of the haplotypes disappears due to deletion and the other haplotype remains one copy or changes to multiple copies. As a result, SNVs from the first outcomes have the potential to contribute imbalance characteristics for the haplotype estimation and hence are possible to be phased. The segments from the other two outcomes are either unable to provide significant evidence to separate the two haplotypes because of comparable allele depths or possess only homozygous SNVs. For our concern, therefore, SNVs in segments of the first outcome are the targets to be phased.

#### Parameters to be estimated

Assume there are *N* CNV segments concerned. Here we aim to estimate the copy numbers of the major *H* and the minor *h* haplotypes in a tumor sample; denote them as *C_H,i_* and *C_h,i_* for segment *i*, 1 ≤ *i* ≤ *N*. Since normal and tumor cells may coexist in the samples, reads from normal and tumor cells may be mixed in sequencing data. There arises a parameter, tumor purity rate *ρ*, to be concerned. The purity is the proportion of tumor cells in a mixed sample.

We can extract different features from the input datasets, and these features would constrain the parameters.

#### Constraints according to allele depths

From the tumor sequencing data, we can calculate for sequencing depth *D_H,i_* and *D_h,i_* for the major and minor haplotype *H* and *h* for each segment *i*, respectively. Moreover, we can estimate the amplification factor *D*; that is, the number of times a single copy of a haplotype is sequenced in the tumor dataset. *D_H,i_* and *D_h,i_* can be computed from the variant call format (VCF) file, as the average across the loci. We adopted K-means clustering to estimate *D*. We normalize the depth of a segment by the depth of the whole tumor data set to calculate the copy numbers initially. Then we choose the number of integers from the rounded minimum copy numbers as the the number of clusters for the K-means. Finally, we pick half of the average depth of the segments in the cluster with the minimum cluster centroid as *D*.

Now, we can constrain *D_H,i_* and *D_h,i_* as

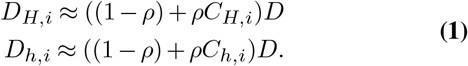

#### Constraints according to segment depth

From the input, we can calculate the average depth *D_i_* for segment *i*, hence, we can constrain the parameters as

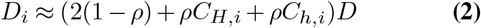

#### Constraints according to depth differences

The average difference *S_i_* can be computed from input between the two haplotypes for segment *i*, and it should be also comparable with the one calculated from *ρ, C_H,i_* and *C_h,i_*, that is

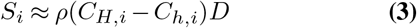

#### Constraints according to allele imbalance

We define Λ_*i*_ as the allele imbalance (AI) for segment *i*, which is a weighted average of the AI values at all the heterogeneous loci in the segment. Assume the segment *i* harbours heterogeneous locus set *K_i_*. Denote λ*_k_* and *w_k_* as the AI value and weight of locus *k*, respectively. Then we have

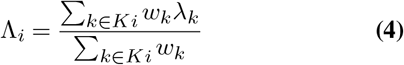

Below we specify how to obtain λ*_k_* and *w_k_*. Denote the allele depths as *d_k,r_* and *d_k,a_* at the *k*-th locus for reference, and alternatives, respectively, then 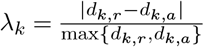.

Under the assumption that the reference and alternative alleles will be sequenced by equal chance if the segment is balanced, i.e., *q_k,r_* = *Pr*(reference is sequenced) = 0.5, the allele depth of a variant should follow a binomial distribution (see Equation Eq. (5)).

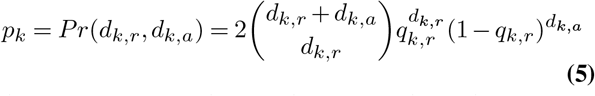

Then we formulate the weight *w_k_* for the *k*-th variant as Equation Eq. (6). Phred quality scores Q are defined as a property which is logarithmically related to the base-calling error probability P

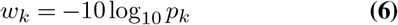

Such a weight has a property that the more imbalanced the allele depths are, the larger the weight will be since the binomial coefficient in *p_k_* will be smaller. The AI Λ*_i_* values should be compatible from these calculated from the parameters; that is,

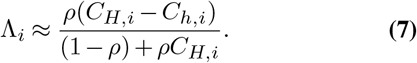

#### Solving the parameters with integer programming

With the aforementioned constraints, we implement an integer programming (IP) to estimate *C_H,i_* and *C_h,i_* for each segment. We replace the approximations by error variables *ϵ*, where we want to minimize the sum of errors.

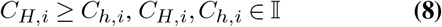

where 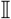 is the integer set.

Combining the constraints Eq. (1), Eq. (2), Eq. (3), Eq. (7) and Eq. (8), we summarize the model in Eq. (9).

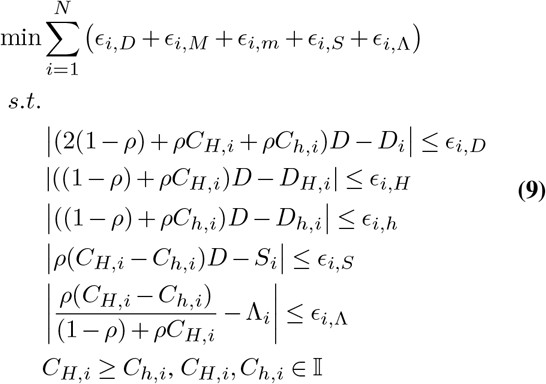

### Haplotype estimation

Having estimated the major and minor copy numbers of segments in pure tumor cells, CNAHap will proceed to perform phasing. With *C_H,i_*’s and *C_h,i_*’s, CNAHap will phase SNVs along the segments where *C_H,i_* > *C_h,i_*. Before phasing, the allele depths of SNVs in each segment will be updated to the allele depth in pure tumor cells. For segment *i*, we first calculate the fractions of major and minor depths (*f_i,H_* and *f_i,h_*) contributed by tumor cells using the purity *ρ* and the major and minor copy numbers, *C_H,i_* and *C_h,i_* (see Equation Eq. (10)).

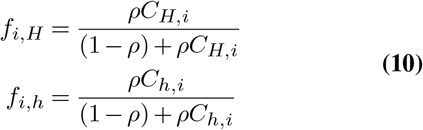

Then we update the allele depths for all SNVs in the segment, multiplying the observed allele depths by the corresponding fraction.

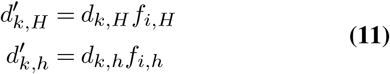

Finally, we perform the phasing for the segment by comparing the updated depths for all SNVs and treating the major alleles of all SNVs as the variants from one haplotype and the minor alleles from the other. Therefore, the two haplotypes *H_i_* and *h_i_* of a segment involving *n* SNVs can be obtained as

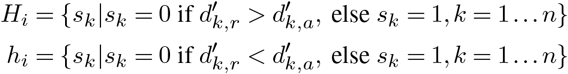

## Results

### Overview of CNAHap

Each individual obtains two copies of chromosomes from parents separately. All genetic markers along the personal genome, such as single nucleotide variants (SNVs), small insertions and deletions (InDels), and short tandem repeats (STRs) should maintain the same across cells, except for somatic mutations occurring in tumor cells. Furthermore, the order and sequence of markers along local regions of chromosomes are the same as these inherited from parents. If copy number variations occur in some genome areas in tumor cells, these regions will gain extra copies. Hence, one or both haplotypes regarding these regions should be duplicated multiple times. Therefore, copies of two haplotypes in a genomic region in tumor cells may become imbalanced, and variants of markers in such a region may have imbalanced sequencing depths, providing evidence for the original haplotypes (Figure 1a).

Here, we produced a computation tool, CNAHap, that adopts the above mentioned principle as core to phase. The skeleton for CNAHap was illustrated in Figure 1b. First, with paired tumor and normal BAM files, CNV caller Accurity (21) is adopted to call the CNV blocks and tumor purity. CNAHap then estimates allele-specific copy numbers for segments of allele imbalance with an integer programming model and filters out those which contain little SNV locus or are allele balanced. Then the phasing algorithm is then performed on each filtered CNV segment along each chromosome. Third, CNAHap outputs the resolved haplotype in VCF format, which benefits subsequent analysis and interpretation. Finally, with auxiliary annotation and downstream analysis scripts, the output of CNAHap can be interactively visualized in CNV: Circos View, CNV: Focal Cluster, and Phased: On Genes, hosted on bio.oviz.org Bio-Oviz (20) (Figure 1c).

### Evaluation of CNAHap on *in silco* data

To evaluate the sensitivity of CNAHap, we invested *in silico* mixtures of sequencing reads from the normal-tumor pair with increased proportions to simulate different tumor purity ratios (20, 50, 80, and 100% tumors). First of all, we evaluated the accuracy (ACC), sensitivity (SE), and specificity (SP) in determining whether to phase segments in Accurity and CNAHap (Figure 2a, Supplementary Table S1a). CNAHap shows a higher accuracy and sensitivity than Accurity. Figure 2b and Supplementary Table S1b display that CNAHap has more correctly called allele-specific CNVs than Accurity. Figure 2c is the histogram plot of phased block length in CNAHap. Despite the purity, the majority (all *>* 93.88%) of phased CNV segments’ block length is larger than 100kbp (Supplementary Table S1c). As illustrates in Figure 2d and Supplementary Table S1d, CNAHap achieves high SNVs phase rates, all larger than 99.98 % regardless of the purities. The number of phased SNVs in purity 1 for samples sim_1, sim_2, and sim_3 is 33740, 32477, and 34436, respectively. With the decrease of the tumor purity, the number of phased SNVs increased. The reason might be that the increase of normal reads in synthetic mixtures adds bias on the CNV segmentation procedure, yielding longer CNV segments qualified for phase. The statistical difference of purity 0.2-0.5 vs. purity 1 is much higher than purity 0.8 vs. 1. In Figure 2e-f and Supplementary Table S1e, we observe that the mean of switch error and mismatch error in purity 1 samples are 0.0270 (SD: 0.0567, Median: 0, IQR: 0-0.0153) and 0.0294 (SD: 0.0471, Median: 0, IQR: 0-0.0441). The switch error and mismatch error on samples with purity 0.2 and 0.5 are significantly higher than purity 1 samples (p-value of SE: 6.7e-16 and 5.8e-05, p-value of mismatch error: < 2.22e-16 and 1.2e-05). In contrast, samples between purity 0.8 and 1 tell no significant difference in error rate. To summarise, our synthetic experiments reveal that as long as the tumor purity larger than 0.5, CNAHap enables producing trustable copy numbers and phase profiles.

**Fig. 2.**
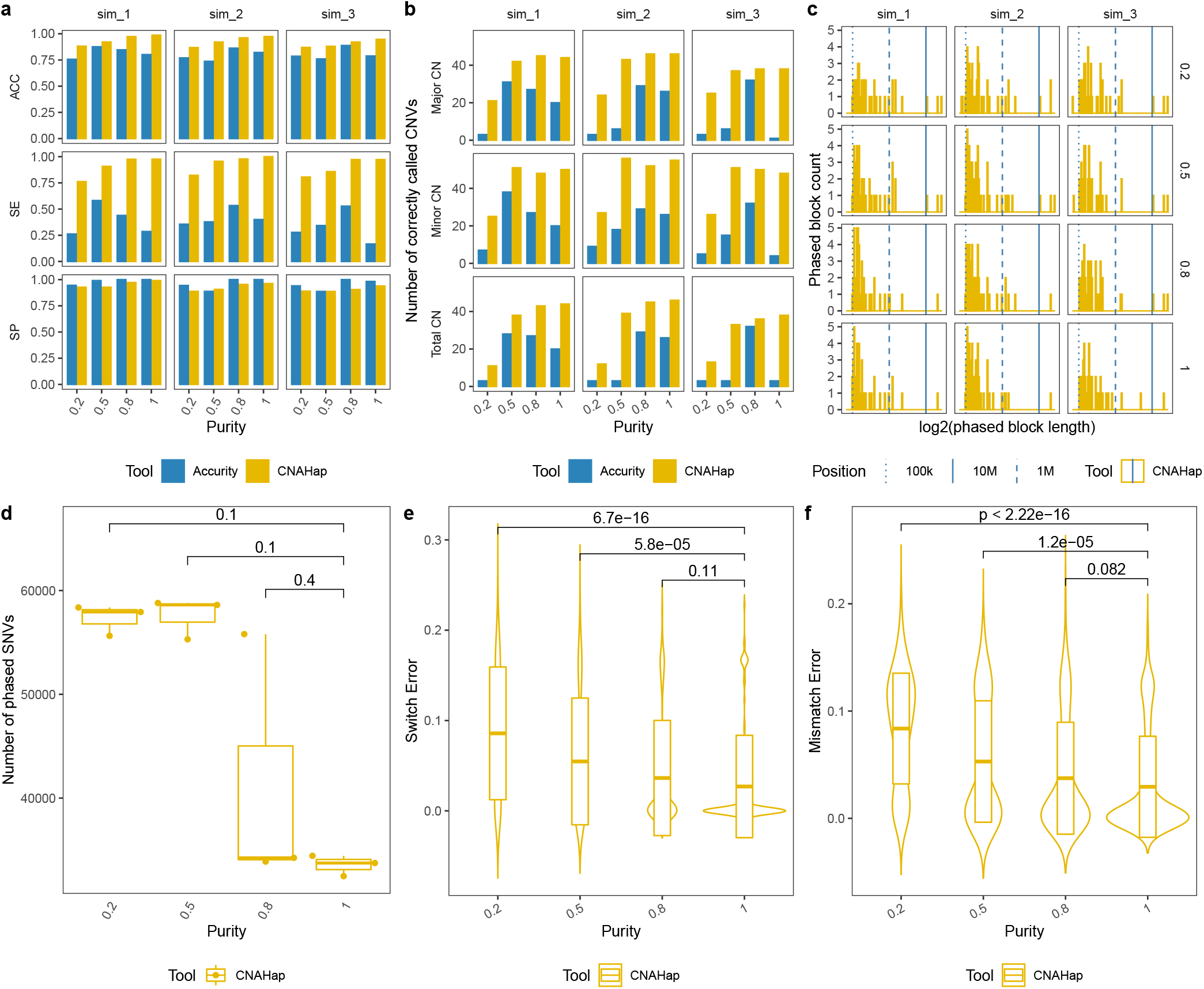
Evaluation of synthetic data among different ploidy. (a) The accuracy (ACC), sensitivity (SE), and specificity (SP) in determining whether to phase CNV segments in Accurity and CNAHap. (b) The number of correctly called CNVs in Accurity and CNAHap. (c) The histogram plot of phased block length in CNAHap. (d) The number of phased SNVs in CNAHap. (e-f) The swith error and mismatch error in CNAHap.

### Case study on a Hepatocellular carcinoma cohort

Hepatocellular carcinoma (HCC) is one of the leading causes of cancer death (24). Sung *et al*. have studied hepatitis B virus (HBV) integration in liver cancer genomes by leveraging the whole-genome sequencing of HCC tumors and adjacent normal tissues (25). In the present case study, we applied CNAHap to reanalyzed the data focusing on putative cancer-related gene amplification with phased haplotypes.

As experimented in *in silico* datasets, CNAHap is sensitive to tumor purity. Thus, we filtered out all samples smaller than or equal to 0.5 (Figure 3a). Then, we selected tumors with prevalent large copy number abberations across the genome (Figure 3b). As a result, 24 HCC samples remained. The circos plot Figure 3c demonstrates the 24 HCC samples are prevalent with copy number gains and allele imbalance across the genome. CNAHap also deciphered the major and minor copy number of each CNV segment. We run GISTIC2 (26), RAIG (22), and RUBIC (23) to check the focal CNV events. As illustrated in Figure3d and Supplementary Table S2a, RUBIC detected one significant (q-value < 0.25) amplification region chr20:25849750-30020750; RAIG detected 10 significantly (q-value < 0.25) amplified region: chr1:15001-563000, chr4:14001-68500, chr5:1547501-1920500, chr5:17635001-17922500, chr8:12046001-12315500, chr8:1923001-2332000, chr10:38769001-38889000, chr12:1-148500, chr14:106785501-107289000, and chr14:19000001-19153000. We abandoned GISTIC2 as it produced lots of focal deletions in contrast with the truth of no deleted segment was called among 24 HCC samples (Figure 3c). Among the 11 focal CNV events, 53 genes were annotated (Supplementary Table S2a), and their total, major, and minor copy number are depicted in the heatmap Figure 3e. We found that 12 genes are previous reported to show focal CNV event in another Chinese HBV associated HCC cohort (27) (focal gains on *DEFB109P1B, FAM138A, FAM138F, FAM66A, LOC100132062, LOC100132287, LOC100133331, OR4F16, OR4F29, OR4F3, OR4F5;* focal loss on *FAM86B2*). (Supplementary Table S1b). Figure 3f demonstrates that focal amplified genes were significantly enriched (p-value < 0.05) in 18 GO pathway and 1 KEGG pathway. Olfactory transaction pathway/olfactory receptor activity (focal gains on *OR4F16, OR4F3, OR4F29, OR4F5, OR11H12*) are recognized as putative drivers of cancer (28). NADH dehydrogenase activities were associated with HCC (29). Kaszak *et al*. reported that cadherin binding associated with HCC (30).

**Fig. 3.**
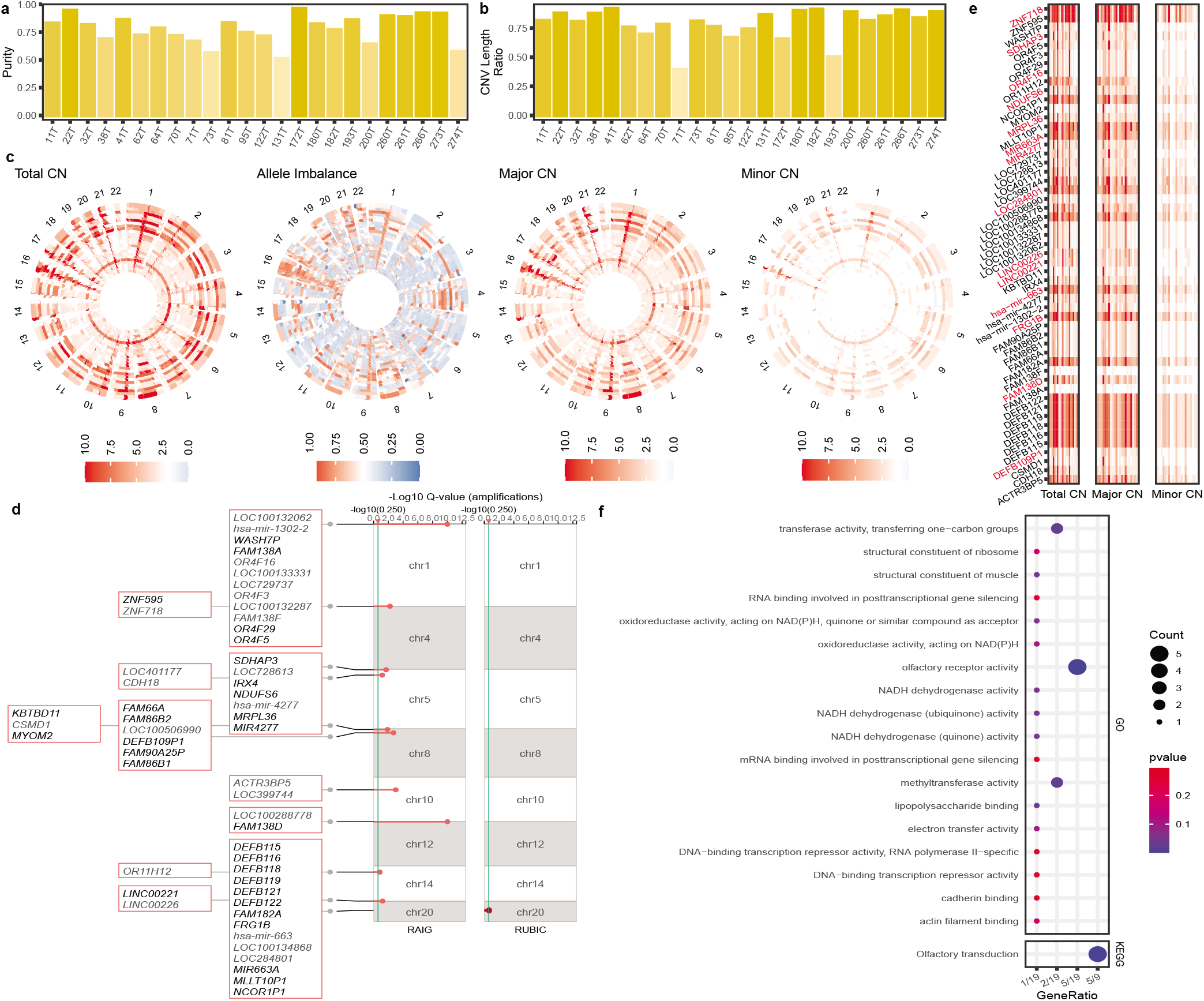
HCC CNV profiles. (a) The estimated purity. (b) The CNV Segments region genomic proportion. (c) The circos plot of total CN, allele imbalance, major CN, minor CN across the genome, one circos layer represents one HCC sample. (d) The focal gains obtained from RAIG (22) and RUBIC (23). (e) The CNV heatmap of focal gain genes. (f) The enriched GO and KEGG pathway of focal gain genes. In (c) and (e), allele imbalance larger than, equal to, and less than 0.5 is annotated in red, white, and blue, respectively. Total CN larger than, equal to, and less than 2 is colored in red, white, and blue, respectively. Major/Minor CN larger than, equal to, and less than 1 is labeled in red, white, and blue, respectively. HCC: Hepatocellular carcinoma, CN: Copy Number.

Figure 4 demonstrates the CNAHap phasing profiles among 24 HCC samples. In Figure 4a, we observe that the phased CNV segments were dominant across the whole genome with the mean of genomic region proportion 78.78 (SD: 13.01, Median: 81.29, IQR: 75.23-87.97)% (Supplementary Table S3a-b). Despite the number of phased CNV segments (Mean: 377.17, SD: 282.34, Median: 288, IQR: 155-578.25) varying across samples, the ratio of phased CNV segments is on average at 49.56 (SD: 09.22, Median: 46.28, IQR: 47.43-55.63)% (Figure 4b, Supplementary Table S3a-b). Figure 4c-d demonstrate that CNAHap generates large phasing blocks. The average N50 and N90 among HCC corhot is around 25M and 7M, respectively. [N50 (Mean: 25,871,765, SD: 19,222,539, Median: 21,537,499, IQR: 9,630,624 - 42,678,249) bp, N90 (Mean: 7,456,541, SD: 6,287,567, Median: 5,764,999, IQR: 2,701,124 - 10,852,249) bp]. The average number of phased SNVs is 1,763,850 (SD: 323,620.1, Median: 1,839,513, IQR: 1,695,264-2,006,223) and the average phase rate is 78.83 (SD: 13.97, Median: 82.26, IQR: 73.72-89.65)%. The long phasing block and high phasing rate are due to the phased CNV events occupying the majority of genome (Figure 4a), and around 93.77 (SD: 3.16, Median: 93.98, IQR: 93.01-95.69)% phased CNV segments are longer than 100k and around 66.78 (SD:10.50, Median: 66:31, IQR: 57.79-76.23)% longer than 1M (Figure 4e, Supplementary Figure S1a-b, Supplementary Table S3a-b), providing extreme long allele imbalance linkage to phase.

**Fig. 4.**
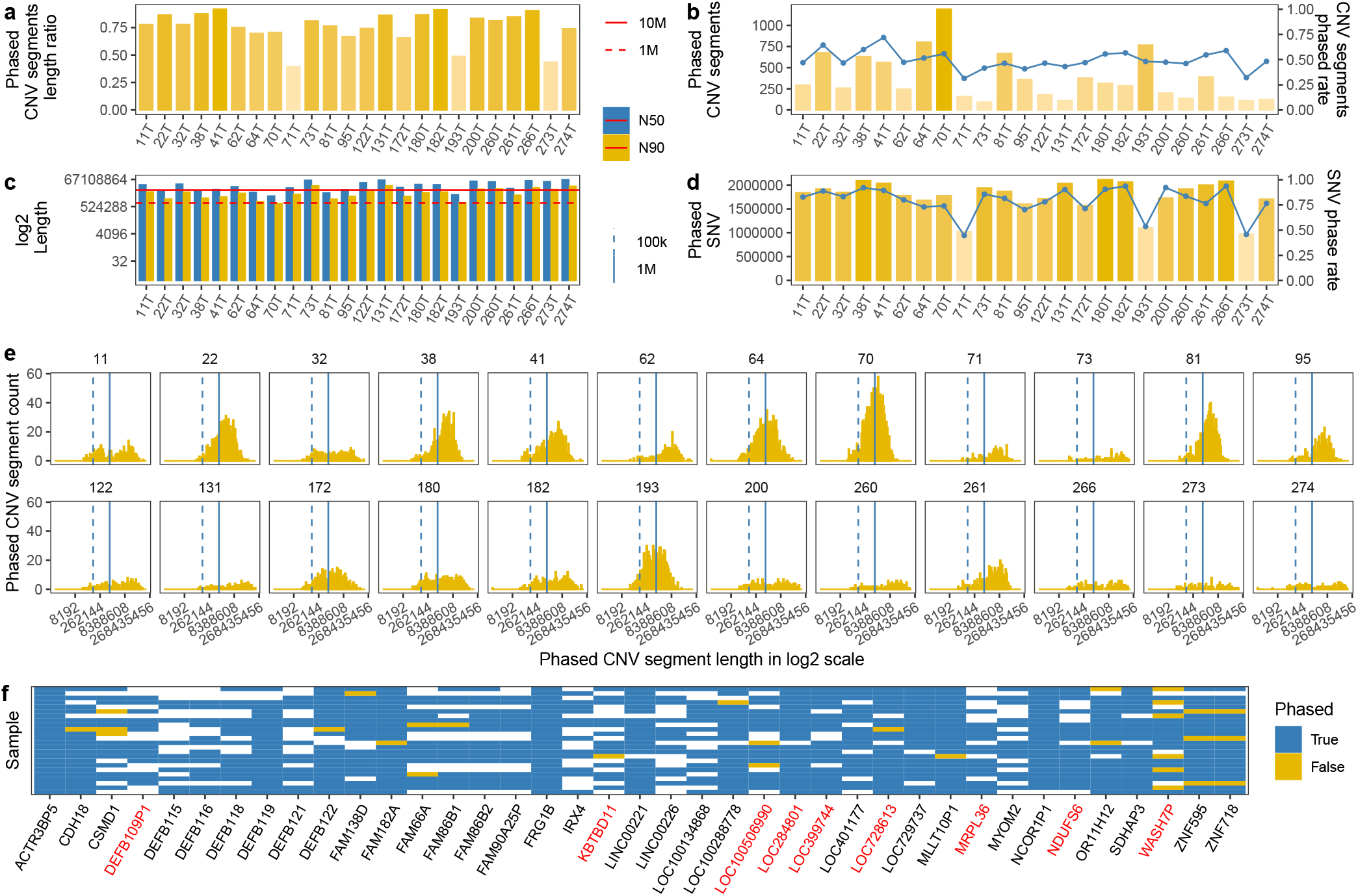
HCC phased profiles among HCC samples. (a) The purity of tumor samples. (b) The number and proportion (blue line) of phased CNV segments. (c) The number and proportion (blue line) of phase SNVs. (d) The N50 and N90 of phased blocks, dashed and solid line indicates the length of 1M bp and 10M bp, respectively. (e) The histogram plot of phased CNV block length. Dashed and solid blue line shows the length of 100k bp and 1M bp, respectively. (f) Overview of phase result of focal gain gene, blue means phased gene, yellow otherwise. White tile indicates there is no SNVs in that gene. Genes colored in red are in enriched GO and KEGG pathway.

Then, we checked the phasing results of focal amplified genes. As demonstrated in Figure 4f, a total of 39 focal gain genes harbor SNV variants. Supplementary Figure S1c illustrates the scatter plot of focal gain gene mutation number and density, we can observe that CNAHap successfully phase genes with larger than 4,000 SNV variants. 23 out of 39 genes are completely phased in the cohort. Among them, LinkRNA *LINC00221* are reported as a potential diagnostic and prognostic biomarker in HCC (31) (Figure 5a). *FAM86B2* also shows focal CNV event in another Chinese HBV associated HCC cohort (27) (Supplementary Figure S2). 15 genes have less than four unphased samples, *WASH7P* has six unphased samples (Figure 5b). The mutation and phasing details for the rest of focal genes can be interactively visualized in web interface “Phased: On Genes” (https://bio.oviz.org/demo-project/analyses/Phased_on_genes, Demo File: “CNAHap_HCC”).

**Fig. 5.**
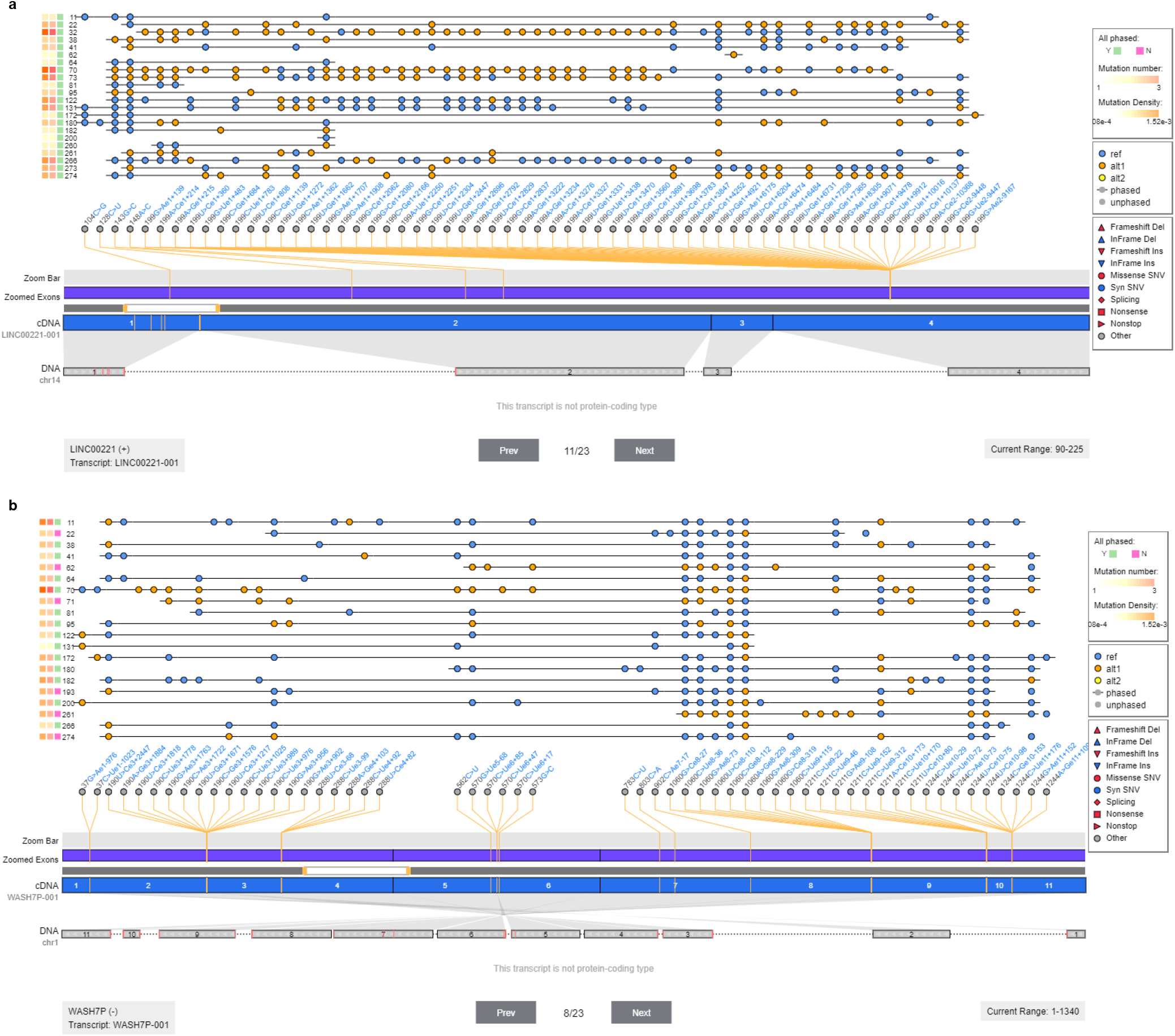
(a-b) Phasing profile on lincRNA *LINC00221* and gene *WASH7P* among the HCC cohort.

### CNAHap online visualization interfaces

The CNV profiles and phasing profiles in text format are nonintuitive for users to perceive the landscape and differences within a patient cohort. Thus, we developed three online web interfaces (CNV: Circos View, CNV: Focal Cluster, and Phased: on Genes) to visualize the output of CNAHap. Table 1 summarises the key features of the provided interfaces. CNV: Circos View, demonstrated as Figure 3a, displays the circos plot of total CN, major CN, minor CN, allele imbalance, phased information, etc., across the patient cohort. CNV: Focal Cluster, showed as Figure 3b, shows the recurrent gains and losses detected by multiple tools, and supports Ensembl (32) annotation. Phased: On Genes, illustrated as Figure 5 and Supplementary Figure S2, displays the mutation detail and phasing profile on genes, supporting transcript (Ensembl (32)) and protein (Pfam (33)) annotation. Generally, we offer the users an editor to upload the CNVHap outputs to the server and adjust the figure display settings. We provide interactive tooltips to show the essential information of a sample, a CNV segment, an SNV variant, and so on, assisting users in seizing potential findings quickly. With one-button clicked, users can download high-quality figures for share or paper publishing. For demonstration, we have uploaded the raw data of Figure 3a, Figure 3b, Figure 5, and Supplementary Figure S2 as demo data set “CNAHap_HCC” in the editor.

## Discussion

Although the heterozygous allelic imbalance from tumor tissue is widely utilized to infer somatic copy number alterations (SCNAs) (21, 34, 35). Collaborating tumor allelic imbalance to phase germline variants has not been broadly adopted. Prepemariy studies on VAF phasing and HATS have established that this data attribute to the assembly of the germline haplotype. However, running these tools requires arduous efforts as VAF phasing provides no accessible source code (18), and HATS necessitates a training process first (19). Thus, we introduce CNAHap, an easy-use tool that leverages imbalance in SNV or InDel alleles in copy number gains region to phase germline haplotype. Like haplotype assembly tools, CNAHap only demands sequencing data and can phase rare and de novo variants. Surpass the assembly-based ones, CNAHap is not constrained by the read length and insert size of particular sequencing protocols, thus yields much greater phasing blocks. CNAHap also calls the allele-specific copy number aberrations in tumor cells.

The allele-specific CNV profiles and phasing profiles in text format are nonintuitive for users to perceive the landscape and differences within a patient cohort. To address this issue, we developed three online web interfaces (CNV: Circos View, CNV: Focal Cluster, and Phased: on Genes) to visualize the output of CNAHap. Equipped with interactive tooltips and editors, users can capture and share potential scientific discoveries without effort.

Noteworthily, some caveats need to be addressed. (1) CNAHap now only phases over the SCNA segments with allele imbalance but does not assign haplotype order in a balanced or diploid genomic region. In other words, CNAHap heavily pivots on the popularity and proportion of imbalanced somatic copy number alterations (SCNAs). Even though Compton *et al*. claimed that large CNV blocks are prevalent across the solid tumor genome (almost 90%) (17), and our HCC case study reported the average SCNA proportion as 78.78 (SD: 13.01, Median: 81.29, IQR: 75.23-87.97)%. There exist near-diploid colorectal cancer (CRC) tumors (36), diploid lymph node metastases (37), diploid endometrioid adenocarcinomas (38), etc. Thus, we recommend fitting the paired normal data to assembly-based tool such as SpecHap (16) simultaneously and combining the phasing results between CNAHap and SpecHap to achieve a complete germline haplotype. (2) Our experiments on synthetic datasets indicate CNAHap is sensitive to tumor purity. The switch error and mismatch error on samples with purity 0.2 and 0.5 are significantly higher than purity 1 samples (p-value of SE: 6.7e-16 and 5.8e-05, p-value of mismatch error: <2.22e-16 and 1.2e-05). In contrast, samples between purities 0.8 and 1 tell no significant difference in error rate. Thus, we advise practicing CNAHap only to tumors with purity larger than 50%. (3) Currently, CNAHap requires SCNA segments as input. We suggest leveraging Patchwork (35) or Accurity (21) to identify SCNA segments first and then using CNAHap to refine the allelic specific copy numbers and the germline haplotypes in regions where demonstrate an imbalanced SNV/InDel allele. (4) In this study, we only validated the efficacy of CNAHap in pair-end sequencing reads. In fact, CNAHap can apply to any normal-tumor pairs notwithstanding the sequencing technologies, as long as users have the CNV segmentation and point mutation VCF file of the normal-tumor pair. (5) CNAHap is unable to recognize the haplotype of somatic mutations. We are considering it as a future enhancement.

## Conclusion

Haplotype phasing is significant in the study of human genetics. The pervasiveness of the large copy number variant segment in solid tumors brings possibilities to resolve long germline phasing blocks utilizing allele imbalance in tumor data. Although there are such studies, none of them provide easy-use software on the premise of availability and usability. Herein, we present a novel method, CNAHap, to determine the copy number in tumor and then phase germline variants in tumor copy number segments with the aid of allele imbalance. We also provide interactive web interfaces to visualize the copy number and phase landscape of CNAHap. On *in silico* datasets, CNAHap demonstrates higher copy number calling accuracy than the benchmark tool and generates long phasing blocks. On a Hepatocellular carcinoma case study, CNAHap successfully generates huge phase blocks with the average N50 and N90 at 25M and 7M, respectively, and find the Olfactory receptor family is recurrent and amplified. In all, our results illustrate the efficacy of CNAHap in determining tumor copy numbers and their long germline haplotypes.

## Acknowledgements

We would like to express sincere gratitude Mr. Yonghan Yu for his valuable assistance and advice.

## Contributions

S.C.L. supervised the project. S.C.L., B.T., W.J. and L.C. discussed the algorithm. B.T. implemented the algorithm and evaluated the method in simulations. L.C. designed and performed the case study. L.C. and W.J. designed the visualization interfaces. Y.W. and H.L. implemented the visualization interfaces in Oviz-Bio. L.C. and B.T. wrote the manuscript. S.C.L. revised the manuscript. All authors read and approved the final manuscript.

## Data availability

HCC data used in this paper can be retrieved under the accession ERP001196 (25). All experiments can be reproduced with the dedicated version of software with default arguments.

## Software availability

CNAHap source code is deployed at https://github.com/bowentan/CNAHap.

## Supplementary Note 1: Supplementary Figures

**Fig. S1.**
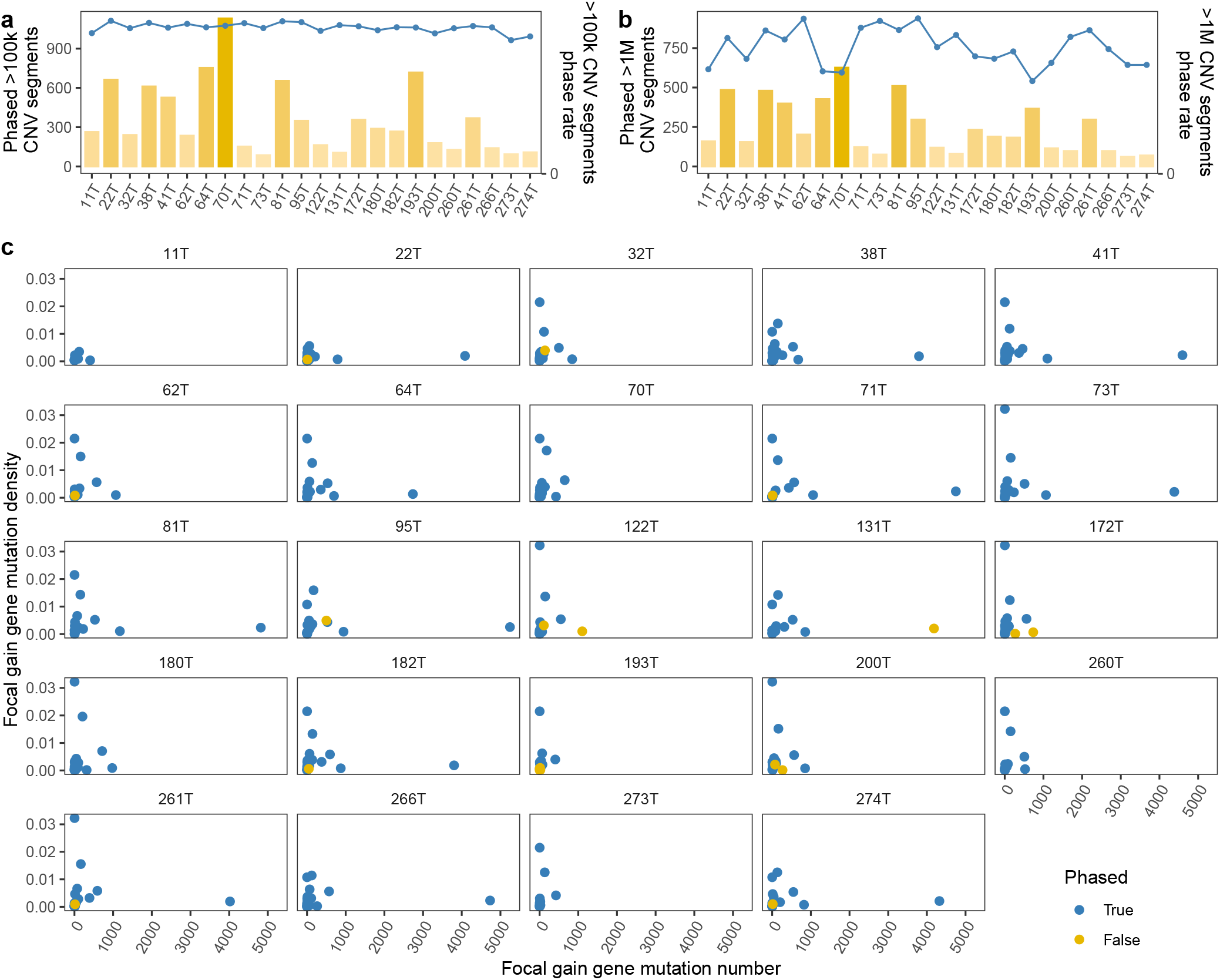
HCC phased profiles among HCC samples. (a) The purity of tumor samples. (b) The number and proportion (blue line) of phased CNV segments. (c) The number and proportion (blue line) of phase SNVs. (d) The N50 and N90 of phased blocks, dashed and solid line indicates the length of 1M bp and 10M bp, respectively. (e) The histogram plot of phased CNV block length. Dashed and solid blue line shows the length of 100k bp and 1M bp, respectively. (c) The scatter plot of focal gain gene mutation number and density, blue means phased gene, yellow otherwise.

**Fig. S2.**
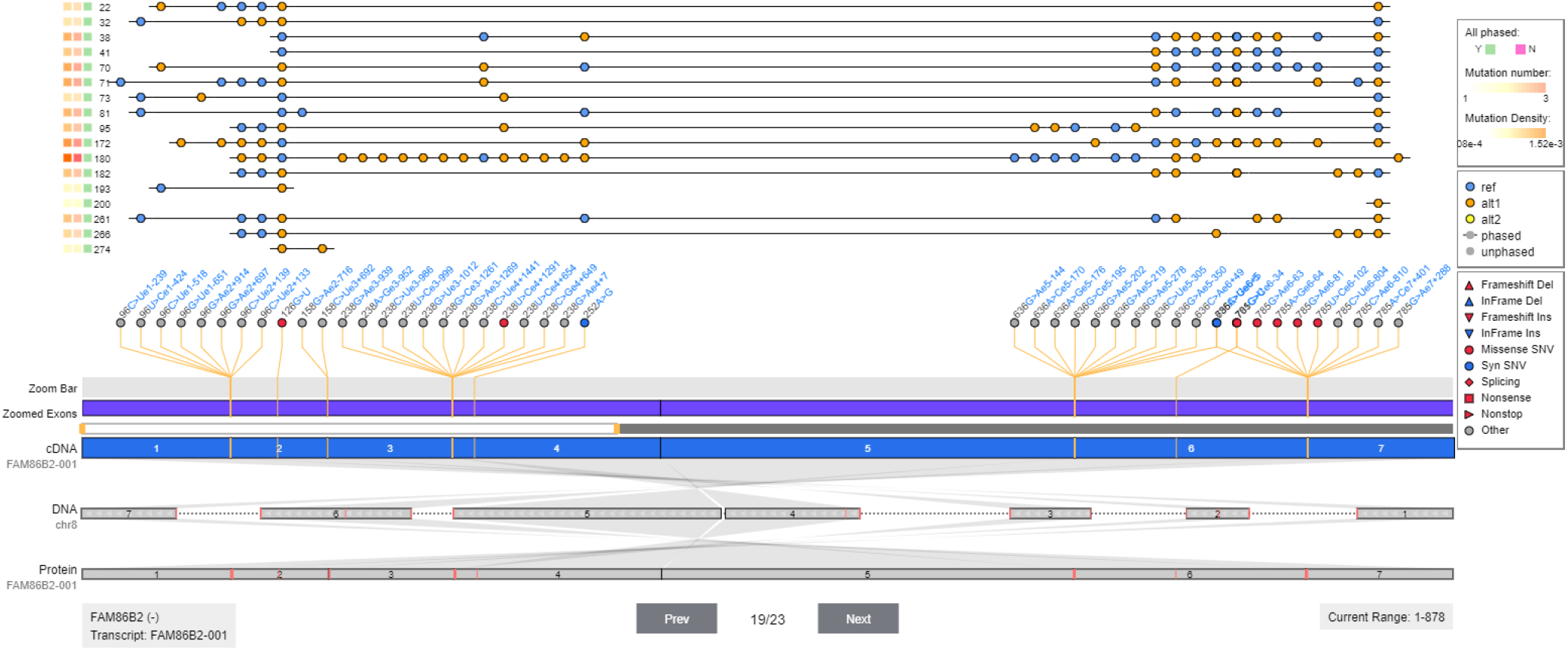
Phasing profile on gene *FAM86B2* among the HCC cohort.

